# Human extinction learning is accelerated by an angiotensin antagonist via ventromedial prefrontal cortex and its connections with basolateral amygdala

**DOI:** 10.1101/512657

**Authors:** Feng Zhou, Yayuan Geng, Fei Xin, Jialin Li, Pan Feng, Congcong Liu, Weihua Zhao, Tingyong Feng, Adam J. Guastella, Richard P. Ebstein, Keith M. Kendrick, Benjamin Becker

## Abstract

Extinction is considered a core mechanism underlying exposure-based therapy in anxiety-related disorders. However, marked impairments in threat extinction learning coupled with impaired neuroplasticity in patients strongly impede the efficacy of exposure-based interventions. Recent translational research suggests a role of the renin-angiotensin (RA) system in both these processes. However, the efficacy of pharmacological modulation of the RA system to enhance threat extinction in humans and the underlying neural mechanisms remain unclear. The present pre-registered, randomized placebo-controlled pharmacological neuroimaging trial demonstrates that pre-extinction administration of the angiotensin II type 1 receptor antagonist losartan accelerated attenuation of the psychophysiological threat response during extinction. On the neural level the acceleration of extinction was accompanied by threat-signal specific enhanced ventromedial prefrontal cortex (vmPFC) activation and its coupling with the basolateral amygdala. Multivoxel pattern analysis and voxel-wise mediation analysis further revealed that that losartan reduced the neural threat expression, particularly in the vmPFC, and confirmed that acceleration of extinction critically involved treatment-induced modulation of vmPFC activation. Overall the results provide the first evidence for a pivotal role of the RA system in extinction learning in humans and suggest that adjunct losartan administration can be leveraged to facilitate the efficacy of extinction-based therapies.

## Introduction

Extinction learning refers to the attenuation of a previously learned defensive response when the threat predictive stimulus is repeatedly encountered in the absence of adverse consequences. Exposure-based interventions capitalize on extinction learning mechanisms to reduce excessive fear in patients with anxiety-related disorders and are considered an efficient therapy in these illnesses [1]. A significant number of patients however does not adequately respond to exposure therapy [2] and impaired extinction processes are considered one key mechanism underpinning the lack of therapeutic efficacy [3]. Anxiety-related disorders are highly prevalent and associated with significant psychosocial impairments and societal costs [4]. As such, innovative strategies to improve the efficacy or shorten the duration of exposure therapies are urgently needed.

Whereas the behavioral and neural pathomechanisms underlying anxiety disorders are increasing understood, translation into efficacious clinical interventions remains inadequate. Notably, the neural mechanisms mediating extinction are extremely well-conserved over the course of evolutionary time; hence, the identification in animal models of receptor targets sensitive to pharmacological modulation that facilitate neural plasticity in pathways supporting extinction learning, can feasibly augment the efficacy of exposure-based interventions [5].

Animal models and human neuroimaging research have demonstrated a crucial role of the infra-limbic cortex (IL), which is homologous to the human ventromedial prefrontal cortex, vmPFC, and its interactions with the amygdala in extinction learning [6-12]. The vmPFC is critically engaged in the reduction of threat expression during extinction [3, 9, 13] and governs the amygdala inhibition of conditioned threat response [8, 10]. Translational models suggest that dysfunctions in amygdala-prefrontal neuroplasticity contribute to extinction-failure in anxiety disorders [14]. Converging evidence from clinical research suggests that anxiety disorders are characterized by deficient extinction, hypoactivation within the vmPFC and attenuated vmPFC-amygdala functional connectivity [6, 8, 11, 13, 15].

Intriguingly, recent evidence suggests that the renin-angiotensin (RA) system, primarily known for its role as a blood pressure and renal absorption regulator, represents a promising target to facilitate extinction [5]. Central angiotensin receptors are densely expressed in limbic and prefrontal brain regions and are critical to changes in neuroplasticity and extinction [16-18]. Initial studies in rodents have demonstrated the potential of pharmacological modulation of RA signaling towards facilitating extinction using the selective competitive angiotensin II type 1 antagonist losartan (LT) [19, 20]. LT is an approved treatment for high blood pressure with an excellent safety record [21, 22]. Notably, initial clinical observations suggested unexpected beneficial effects on memory and anxiety-symptomatology [5, 21]. Taken together, accumulating evidence suggests that LT-modulation of central RA signaling represents a promising target to enhance extinction learning.

Against this background, we conducted a pre-registered randomized placebo-controlled pharmacological experiment to determine whether targeting the RA system can facilitate extinction in humans. To uncover the underlying neural mechanisms functional magnetic resonance imaging (fMRI) and psychophysiological threat responses (skin conductance, SCR) were simultaneously acquired. The specific goal was to determine the potential of LT (50mg, single-dose, p.o.) as a therapeutic candidate for the clinical augmentation of extinction learning. Based on previous translational research we expected that LT would (1) accelerate attenuation of the psychophysiological threat responses, and that enhanced extinction would be mediated by two neural processes: (2) increased activation in the vmPFC and attenuation of its threat expression in the context of (3) stronger functional interaction of the vmPFC with the amygdala.

## Materials and Methods

### Participants and experimental protocols

Seventy healthy males underwent a validated Pavlovian threat acquisition and extinction procedure with simultaneous fMRI and SCR acquisition. To reduce variance related to hormonal and menstrual-cycle-associated variation in extinction [23-25] only male participants were included in the present proof-of-concept experiment [for similar approach see ref. 26]. Due to technical issues (SCR recording failure n = 3) or absence of threat acquisition (n = 8) data from 11 subjects were excluded leading to n = 30 LT- and n = 29 PLC-treated subjects for the evaluation of the primary study hypotheses. For exclusion criteria and a description of the study sample see **Supplementary methods.**

The experiment consisted of three sequential stages: (1) acquisition, (2) treatment administration and, (3) extinction. 20-min after acquisition, participants were administered either a single 50mg (p.o.) dose of the selective, competitive angiotensin II type 1 receptor antagonist losartan (LT, Cozaar; Merck, USA) or placebo (PLC), packed in identical capsules. Consistent with the pharmacodynamic profile of LT [27] extinction was observed 90min post-treatment. Although previous studies reported no effects of single-dose LT on cardiovascular activity or mood [18, 27, 28], these indices were monitored to primarily control for unspecific effects of treatment and not focused on behavior per se. The experimental time-line is provided in **Fig. 1a** (for the pharmacodynamic profile and assessment of confounders see **Supplementary Methods**).

**Fig 1.**
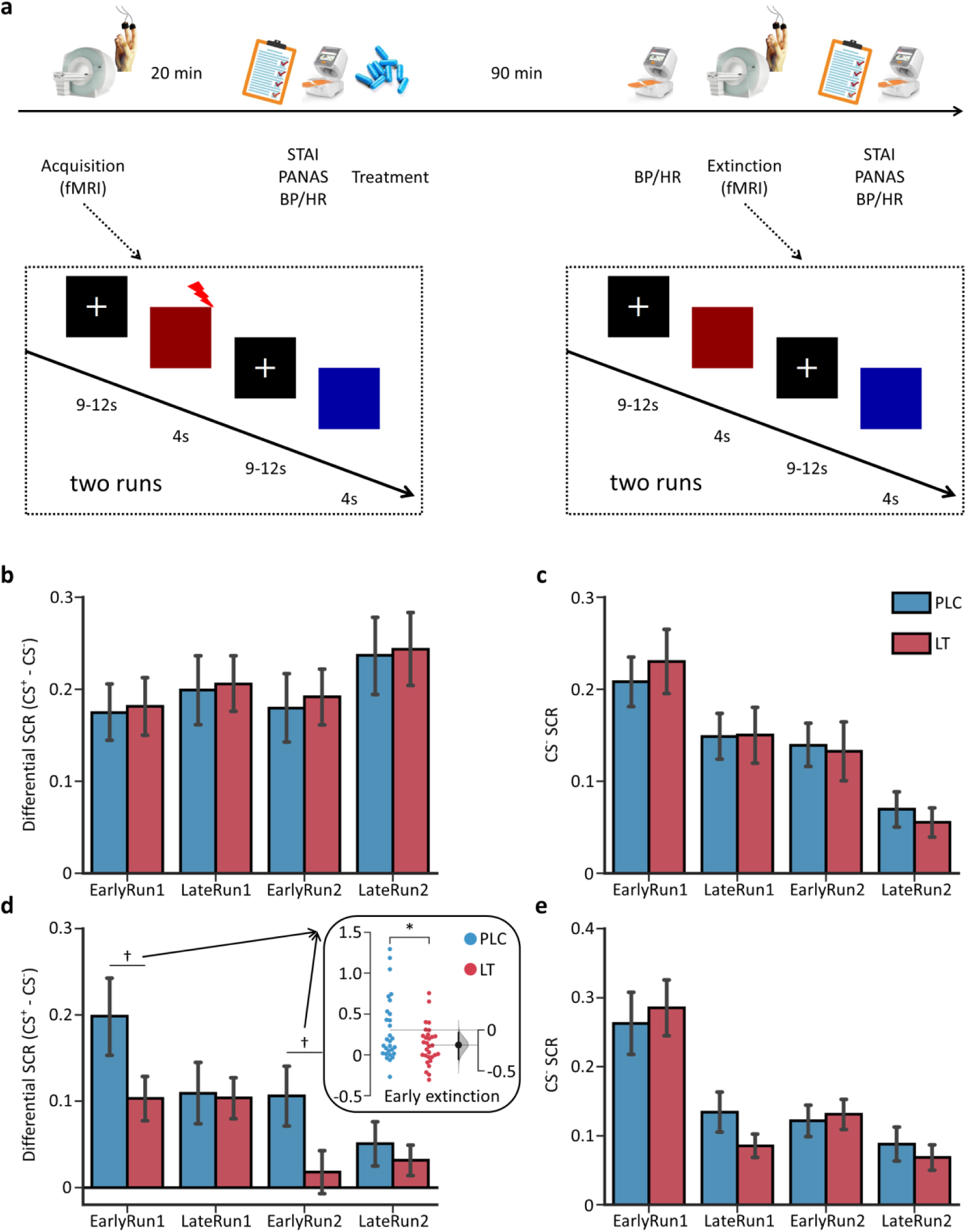
Experimental timeline and losartan effects on psychophysiological threat responses. (a) Experimental timeline and schematic synopsis of fMRI tasks. (b) Psychophysiological threat responses (CS^+^ - CS^-^) during acquisition demonstrating successful CS discrimination with enhanced SCR to the CS^+^ relative to the CS^-^ in both groups. (c) Mean SCR for CS^-^ presentations during acquisition. (d) Psychophysiological threat responses (CS^+^ - CS^-^) during extinction learning. Psychophysiological threat responses during early extinction (across two runs) are presented in the inset. (e) Mean SCR for CS^-^ presentations during extinction learning. †*P* < 0.05, one-tailed; \**P* < 0.05, two-tailed, error bars represent standard errors. The filled curve indicates the null-hypothesis distribution of the difference of means (Δ) and the 95% confidence interval of Δ is illustrated by the black line.

The study was approved by the local institutional ethics committee and adhered to the Declaration of Helsinki and was a pre-registered (ClinicalTrials.gov, NCT03396523).

### Experimental Paradigm

During the acquisition stage, participants were repeatedly presented with two different colored squares, the CS (conditioned stimulus). One CS (CS^+^, 4s) was pseudo-randomly paired with a mild electric shock (US, 2ms) with 43% contingency, whereas the other CS (CS^-^, 4s) was never paired with a US. Acquisition was followed by extinction, where the same cues were presented without US (**Supplementary methods**). To enhance threat memory acquisition as well as to increase the statistical power to determine treatment effects, both learning phases included two subsequent runs of the task [for a similar approach see ref. 29]). Prior to each run, subjects were informed that “the experimental runs are independent and you may or may not receive the electric shock” thus subjects were unable to predcit the presence or absence of the US at the beginning of the respective run.

### Skin Conductance Response Analysis

Skin conductance responses were computed as done in previous studies [9, 30-32]. Psychophysiological threat responses were defined as baseline-corrected CS^+^ by subtracting the mean responses to the CS^-^ (see **Supplementary Methods**). Treatment effects were determined employing a phase (early, late) × run (run1, run2) × treatment (LT, PLC) 3-way mixed ANOVA with psychophysiological threat responses as dependent variable.

### MRI Acquisition and Analysis

MRI data were acquired using a Siemens TRIO 3-Tesla system with a 12-channel head coil. Functional time-series were processed using SPM12 (Statistical Parametric Mapping, https://www.fil.ion.ucl.ac.uk/spm/software/spm12/). On the first-level, the CS^+^ and CS^-^ stimuli were modeled and condition-specific regressors for early (first half) and late (second half) of extinction were defined (details see **Supplementary Methods**).

### Whole-Brain Analyses

Effects of LT on extinction were assessed using a whole-brain phase (early, late) × run (run1, run2) × treatment (LT, PLC) 3-way mixed ANOVA with the CS^+^ > CS^-^contrasts as dependent variable. Significant interaction effects were further disentangled by two independent post hoc approaches to warrant both high regional-specificity (whole-brain voxel-wise post hoc t-tests) and high robustness (leave-one-subject-out cross-validation (LOSO-CV) procedure [33]). Group-level analyses (including LOSO-CV) were conducted using FSL Randomise (FMRIB Software Library, http://fsl.fmrib.ox.ac.uk/fsl/fslwiki/) with permutation-based inferences (10,000 permutations). Significant clusters were determined using a height threshold of *P* < 0.001 (two-tailed) and an extent threshold of *P* < 0.05 (two-tailed) with cluster-based family-wise error (FWE) correction.

### Region-of-interest - vmPFC contributions during early extinction

Due to the critical role of the vmPFC in extinction [8, 34-37] and our *a priori* regional hypothesis, we explored LT effects on stimulus-specific vmPFC activation during early extinction. To this end, activity estimates were extracted from an anatomically-defined vmPFC region of interest (ROI) (**Supplementary Methods**) and subjected to a stimulus (CS^+^, CS^-^) × run (run1, run2) × treatment (LT, PLC) 3-way mixed ANOVA.

### Neural Threat Expression - Multi-Voxel Pattern Analysis (MVPA)

Following Reddan, et al. [36], a neural pattern of threat was developed to differentiate CS^+^ versus CS^-^ (trained on the acquisition data) and subsequently applied to early extinction activation to determine treatment effects on the neural threat expression (**Supplementary Methods**).

### Voxel-wise Mediation Analyses

To determine whether treatment effects on vmPFC activation (CS^+^ > CS^-^) during early extinction critically contributed to the accelerated attenuation of the psychophysiological threat responses (SCR, CS^+^ > CS^-^), voxel-wise mediation analyses were conducted (Mediation Toolbox, https://github.com/canlab/MediationToolbox) [38, 39]. Mediation effects were inferred using bootstrapping (10,000 replacements) and false discovery rate (FDR) correction.

### Network-level Effects - Functional Connectivity Analysis

Given that both human and animal studies strongly implicate vmPFC-mediated inhibition of the amygdala as a key extinction mechanism [10-12, 34], a functional connectivity analysis [40] was employed to determine treatment effects on the vmPFC-amygdala coupling during early extinction. We hypothesized that LT-induced extinction enhancement would be accompanied by stronger functional interaction between these regions (one-sided).

## Results

### Participants

Consistent with previous studies [18, 27, 28], no effects of drug or placebo on cardiovascular and affective indices were observed, which together with the chance level guesses for treatment, argues against unspecific confounding effects of treatment (**Supplementary Table 1**). During the pre-treatment acquisition phase, both groups exhibited successful threat acquisition on the psychophysiological (**Figs. 1b,c**) as well as neural level (**Supplementary results and supplementary Fig. 1**). Importantly, two-sample t-tests did not reveal between-group activation differences during this stage.

### Reduced Psychophysiological Threat Responses during Early Extinction

A mixed ANOVA model using the psychophysiological threat response during extinction learning as a dependent variable, demonstrated a significant main effect of run (*F*_(1, 57)_ = 17.063, *P* < 0.001, partial η^2^ = 0.230; η^2^ indicates effect size in terms of eta squared) reflecting decreased psychophysiological threat responses in run2 compared to run1 and a marginally significant main effect of phase (*F*_(1, 57)_ = 3.387, *P* = 0.071, partial η^2^ = 0.056) reflecting decreased psychophysiological threat responses during late extinction. Together these results demonstrated successful extinction learning. Moreover, a significant treatment × phase interaction effect (*F*_(1, 57)_ = 5.017, *P* = 0.029, partial η^2^ = 0.081) was observed, with post hoc two-sample t-tests indicating that relative to the PLC group, the LT group exhibited decreased psychophysiological threat responses during early extinction learning across both runs (*t*_(57)_ = −2.179, *P* = 0.034, d = −0.567, d indicates effect size in terms of Cohen’s d, see insert Fig. **in Fig. 1d**), suggesting accelerated extinction learning. Exploratory run-specific analyses confirmed that LT-treatment enhanced early extinction learning during both initial as well as repeated extinction learning (run1 *t*_(57)_ = −1.805, *P* = 0.038, d = −0.470; run2 *t*_(57)_= −2.012, *P* = 0.024, d = −0.525; two-sample t-tests comparing the treatment groups, one-tailed, **Fig. 1d**). No further significant main or interaction effects were observed on the psychophysiological threat responses (all *P*s > 0.4). Nor were any effects of treatment observed on the safety signal (CS^-^) per se (all *P*s > 0.15, **Fig. 1e**) confirming the threat-specific effects of LT and the absence of unspecific treatment effects on the SCR psychophysiological signal.

### Increased Threat-specific vmPFC Engagement during Early Extinction

Whole-brain analysis revealed a significant treatment × phase interaction effect on extinction-related neural activity (CS^+^ > CS^-^) in the vmPFC (peak MNI_xyz_ = [-3, 27,-12], *F*_(1,57)_ = 23.582, *P*_clusterFWE_ = 0.011, k = 187) (**Fig. 2a**). Regional-specificity of treatment effects was demonstrated by voxel-wise whole-brain post-hoc comparisons demonstrating increased vmPFC activity following LT relative to PLC during early (peak MNI_xyz_ = [6, 48, −3], *t*_(57)_ = 4.505, *P*_clusterFWE_ = 0.013, two-tailed, k = 139), but not late extinction (**Fig. 2b**). Results from the LOSO-CV procedure further demonstrated that LT **-** relative to PLC **-** increased vmPFC activation during early (*t*_(57)_ = 3.417, *P* = 0.001, d = 0.890), but not late (*t*_(57)_ = −1.556, *P* = 0.125, d = −0.405), extinction (**Fig. 2c**, **Fig. 2d** displays an overlay of all leave-one-subject-out-ROIs).

**Fig 2.**
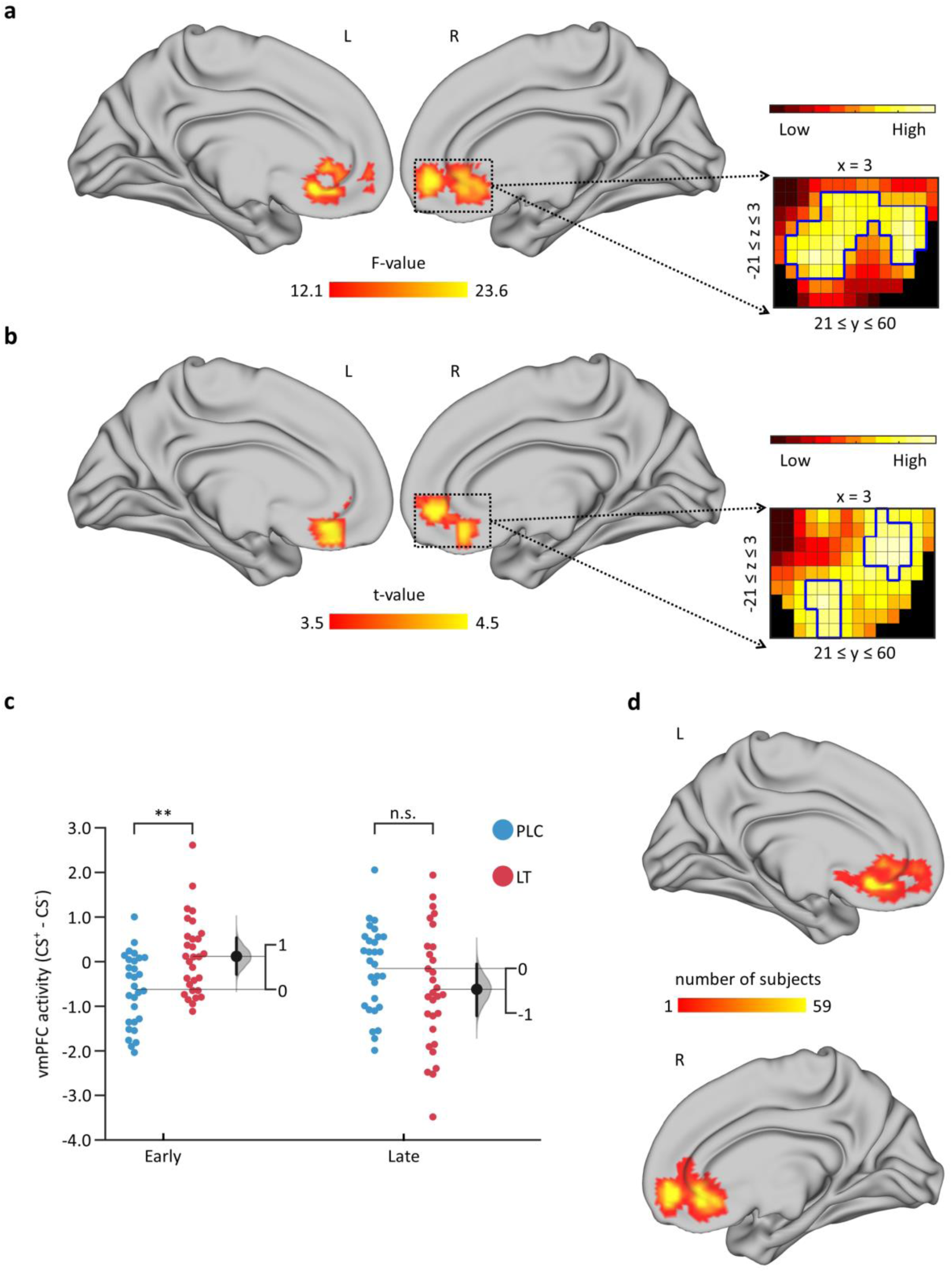
Losartan treatment effects on brain activity (CS^+^ - CS^-^) during extinction learning. (a) vmPFC activity showed significant treatment by phase interaction effect. (b) Losartan specifically increased vmPFC activity during early extinction learning. (c) Mean vmPFC activity (CS^+^ - CS^-^) extracted from the ROIs depicted in (d) showed that losartan increased vmPFC activity during early, but not late, extinction learning. (d) Overlay of all 59 leave-one-subject-out (LOSO) cross-validation (CV) ROIs. All ROIs were created leaving out one subject at the group-level statistic (cluster-level family-wise error (FWE)-corrected). Statistical images were thresholded at *P* < 0.05 (two-tailed), cluster-level FWE-corrected with a cluster-forming threshold of *P* < 0.001 (two-tailed). Examples of unthresholded patterns are presented in the insets; small squares indicate voxel statistical weight; red-outlined squares indicate significance at *P*_clusterFWE_ < 0.05. n.s. represents not significant; \**P* < 0.05. The filled curve indicates the null-hypothesis distribution of the difference of means (Δ) and the 95% confidence interval of Δ is illustrated by the black line. vmPFC, ventromedial prefrontal cortex. vmPFC, ventromedial prefrontal cortex.

Further exploring vmPFC contributions during early extinction by means of extraction of beta estimates, revealed significant main effects of stimulus type and run, as well as a significant treatment × stimulus type interaction effect (details see **Supplementary Results**). Exploratory post-hoc two-sample t-tests demonstrated that LT enhanced threat-specific vmPFC reactivity during both the initial (run1 *t*_(57)_ = 2.141, *P* = 0.037, d = 0.557) and the repeated extinction (run2 *t*_(57)_ = 2.126, *P* = 0.038, d = 0.554), in the absence of effects on the safety signal (CS^-^, *P*s > 0.29) (**Fig. 3a**).

**Fig 3.**
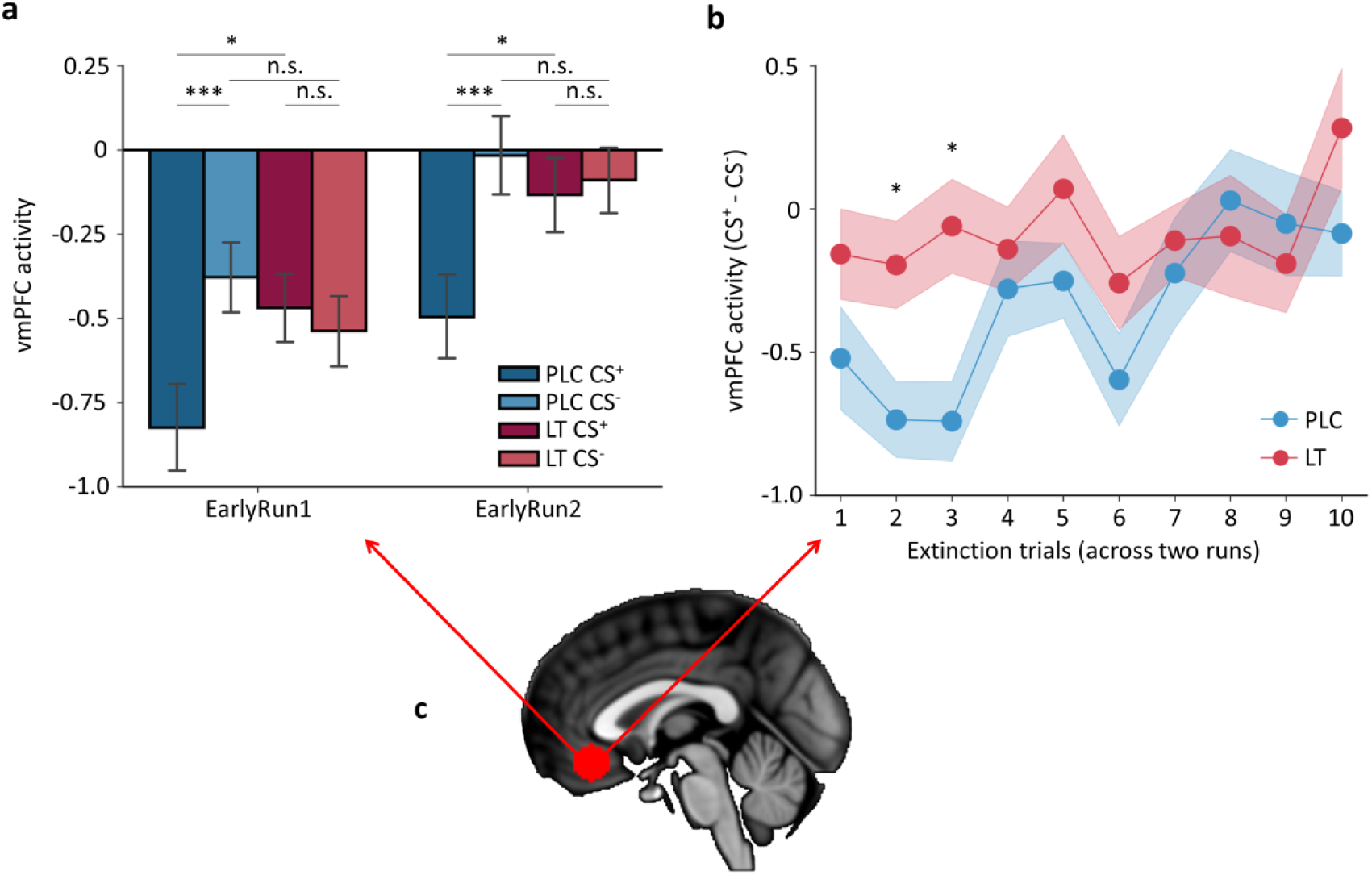
Losartan treatment specifically increased vmPFC activity to threat stimulus (CS^+^) during early extinction learning. (a) Losartan increased vmPFC activity to CS^+^, but not CS^-^, in the early extinction phase in both runs. n.s. represents not significant; \**P* < 0.05; and \*\*\**P* < 0.001, error bars represent standard errors. (b) Single trial analysis confirmed that losartan increased vmPFC activity to CS^+^ in early trials during extinction learning. \**q* < 0.05, FDR corrected. Data are represented as group mean ± SEM. (c) vmPFC ROI. vmPFC, ventromedial prefrontal cortex.

Consistent with our hypothesis on accelerated extinction by LT, an exploratory single trial analysis (**Supplementary Methods**) revealed that LT specifically increased vmPFC activation during initial trials of re-exposure to the threat stimulus (*q* < 0.05, FDR corrected) (**Fig. 3b**).

### Reduced Neural Threat Expression during Early Extinction

The threat-predictive pattern reliably evoked neural threat reactivity during acquisition (comparable to [ref. 36], see **Supplementary Results and supplementary Figs. 2a,b**). Applying the threat-predictive pattern to early extinction activation (CS^+^ > CS^-^) using a LOSO-CV procedure, demonstrated that first, in the entire sample higher neural threat expression was associated with stronger psychophysiological threat reactivity (*r*_(57)_ = 0.571, *P* < 0.001) and confirmed functional relevance of the neural expression of threat [36]. Second, relative to PLC, LT significantly decreased the magnitude of the threat-predictive pattern expression (*t*_(57)_ = −2.091, *P* = 0.041, d = −0.544, two-sample t-test), confirming attenuated neural threat expression during early extinction. Based on our *a priori* regional hypothesis, and the key role of the vmPFC in extinction [3, 8, 9, 13], a vmPFC-focused partial threat expression analysis was conducted.In concordance with the whole-brain results, LT significantly attenuated the vmPFC partial threat pattern expression (CS^+^ > CS^-^) during early extinction (*t*_(57)_ = −3.410, *P* = 0.001, d = −0.888) (see **Supplementary Results** and **supplementary Fig. 2c**).

### vmPFC Activation Drives LT-induced Accelerated Extinction

The voxel-wise mediation analysis aimed at further determining the relationship between treatment, psychophysiological threat attenuation and the underlying neural basis. Conjunction effects (paths a, b and a × b) were observed in a vmPFC cluster (peak MNI_xyz_ = [-3, 45, −15], Z = −3.692, *q* < 0.05, FDR-SVC corrected in the anatomical vmPFC, k = 138), demonstrating that LT increased vmPFC activation (path a), while activation in this region was associated with stronger suppression of psychophysiological threat independent of treatment (path b). Importantly the a × b mediation effect reached significance, indicating that vmPFC activation critically mediated the effects of LT on extinction acceleration (**Fig. 4a**, details see **Supplementary Results**). To test the robustness and visualize the mediation effect, an independent vmPFC-focused mediation analysis was conducted which confirmed the critical contribution of the vmPFC (**Fig. 4B**; a × b effect, bootstrapped *P* value = 0.016; each path is shown in **Fig. 4C**).

**Fig 4.**
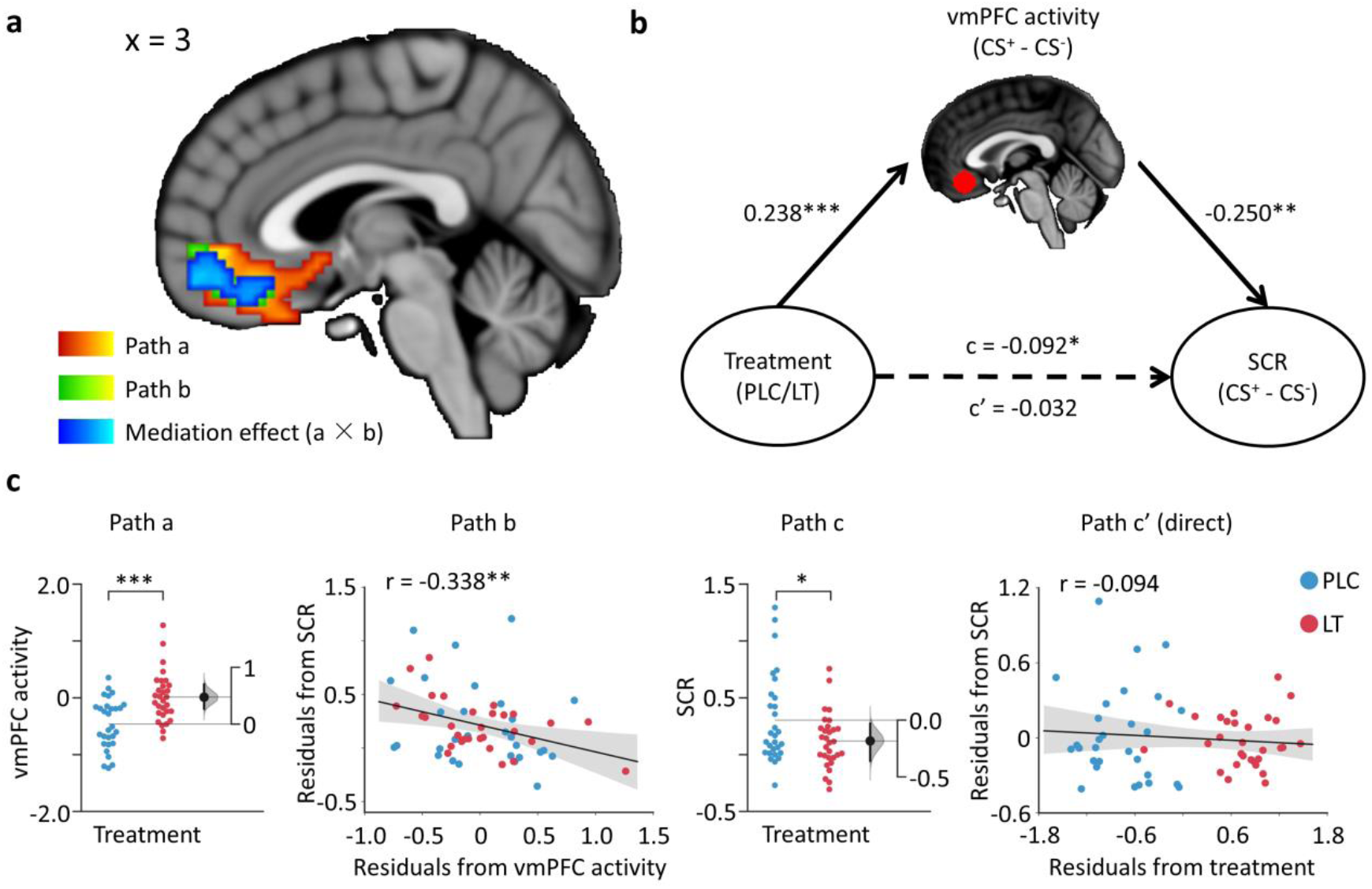
vmPFC activity mediated losartan treatment effect on accelerated extinction learning. (a) Sagittal slice showing regions whose activity increased response to the losartan treatment in yellow (path a), regions whose activity significant negative correlated with psychophysiological threat responses while controlling for the treatment effect in green (path b), and regions whose activity showed significant mediation (a × b) effect in blue. All images were thresholded at *q* < 0.05, FDR corrected within the vmPFC mask. (b) Mediation path diagram with the brain activity in the vmPFC ROI. (c) Examples of each path in the mediation path diagram. \**P* < 0.05; \*\**P* < 0.01; \*\*\**P* < 0.001. The filled curve indicates the null-hypothesis distribution of the difference of means (Δ) and the 95% confidence interval of Δ is illustrated by the black line. vmPFC, ventromedial prefrontal cortex.

### Enhanced vmPFC-Amygdala Coupling

During early extinction LT-induced enhanced functional coupling between the vmPFC and the right amygdala (peak MNI_xyz_ = [24, −3, −24], *t*_(57)_ = 3.557, *q* < 0.05, one-tailed, FDR-SVC corrected in the amygdala, k = 13. This was cytoarchitectonically [41] mapped to the basolateral amygdala BLA, **Fig. 5a**). Subsequent examination of stimulus-specific connectivity estimates from the vmPFC-bilateral BLA pathway, confirmed specific effects of LT on threat signal (CS^+^) processing (*t*_(57)_ = 2.147, *P* = 0.036, d = 0.559) (**Fig. 5b**).

**Fig 5.**
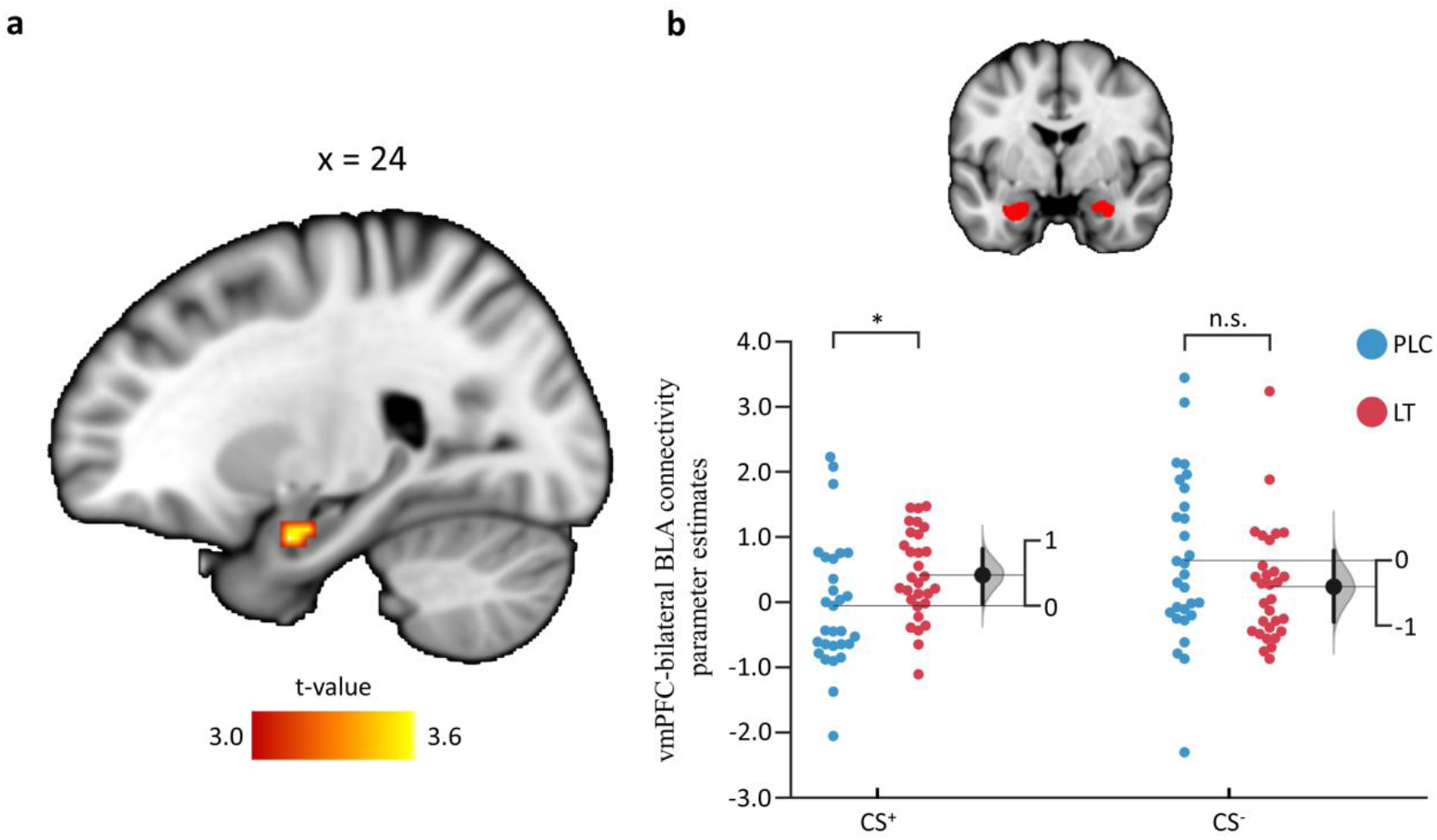
Losartan treatment effect on vmPFC-amygdala functional coupling. (a) Sagittal slice showing that losartan increased functional connectivity between vmPFC and BLA. The image was thresholded at *q* < 0.05, FDR corrected within the amygdala mask. (b) Extracted gPPI parametric estimates in the bilateral BLA showed that losartan treatment specifically enhanced vmPFC-bilateral BLA functional pathway during fear-associated stimulus presentation. n.s. represents not significant; \**P* < 0.05. The filled curve indicates the null-hypothesis distribution of the difference of means (Δ) and the 95% confidence interval of Δ is illustrated by the black line. vmPFC, ventromedial prefrontal cortex; BLA, basolateral amygdala.

## Discussion

The present study demonstrated that LT treatment accelerated the attenuation of a previously acquired psychophysiological threat response, indicating its potential to facilitate threat extinction learning in humans. During early extinction, the acceleration was was critically mediated by enhanced threat-signal specific vmPFC activity and stronger functional coupling of the vmPFC with the BLA. These findings were further paralleled by a pattern classification approach showing that LT-treatment accelerated attenuation of the neural threat expression, particularly in the vmPFC. Overall, the present findings provide first evidence for an important contribution of the RA system to fear extinction in humans and the potential of LT to accelerate extinction through effects on the vmPFC and its inhibitory connections with the BLA.

In humans, successful extinction is accompanied by decreased psychophysiological threat reactivity and concomitantly increased vmPFC activation in response to the threat signal [35, 37, 42]. In the present study LT-treatment reduced the psychophysiological threat responses and selectively enhanced vmPFC activation in response to the threat signal (CS^+^) during early extinction, indicating its potential to accelerate extinction learning in humans. Moreover, the acceleration effects were found not only during the initial extinction learning (i.e., run1), but also in the following “new” learning process (for extinction run2, participants were told that the two extinction runs were independent, and thus they could not predict the absence of the US during the extinction run2). The findings resemble previously observed LT-enhanced extinction learning in rodents [19, 20] and further confirm the important contribution of the vmPFC to successful extinction. Exploring the stimulus-specific effects of LT-treatment on vmPFC activation revealed that reactivity to the safety signal (CS^-^) remained unaffected. Decoding the temporal pattern of LT-effects further suggested that LT specifically attenuated vmPFC reactivity during early re-exposure towards the previously conditioned threat signal. Non-pharmacological stimulation of the vmPFC homologous infra-limbic cortex accelerates extinction learning in rodents [43-45], whereas inactivation or lesion of this region critically impede threat reduction during extinction ([for reviews see refs. 8, 13]). Compatible with the present findings, previous studies demonstrated that non-pharmacological stimulation of the vmPFC can enhance early extinction learning in humans [46], albeit unspecific effects on CS^-^ reactivity have also been reported [47].

Converging evidence from different research focuses suggests that the vmPFC, or the homologous IL in rodents, critically contributes to the reduction of threat expression during extinction learning [3, 8, 13, 34] and regulates amygdala output to inhibit the conditioned threat response [8, 10]. Consonant with the proposed contribution of the vmPFC to extinction learning, LT-attenuated psychophysiological threat responses during early extinction were accompanied by an attenuated neural threat expression, particularly in the anatomically defined vmPFC. Compatible with previous studies that showed critical contributions of the vmPFC to extinction enhancement [43-45] as well as associations between activity in this region and psychophysiological threat reactivity during extinction [48], an additional mediation analysis was carried out. It revealed that higher vmPFC activation was associated with stronger suppression of the psychophysiological threat response. Notably, LT-facilitated suppression of the psychophysiological threat response crucially involved enhanced vmPFC activation (for convergent mediation effects of vmPFC threat expression see **Supplementary Methods and Results**) further emphasizing the key role of this region in extinction enhancement.

On the network level, LT-accelerated threat reduction during early extinction was paralleled by stronger functional communication between the vmPFC and the amygdala, specifically the basolateral subregion. Previous lesion studies in humans demonstrated a critical role of the BLA in threat processing [49] and of the vmPFC in inhibiting amygdala threat responses by exerting top-down control over this region [50]. Animal models further confirmed the importance of pathway-specific neuroplastic changes in the vmPFC-amygdala circuitry during extinction memory formation [10-12] and suggest that vmPFC inputs to the amygdala, instruct threat memory formation and/or gate the expression of conditioned threat [10] during early extinction [12]. The present findings of CS^+^-specific increased vmPFC-BLA functional connectivity following LT-treatment likely reflects an important modulatory role of angiotensin signaling on vmPFC regulation of the amygdala. Previously, animal models demonstrated that stimulation of vmPFC inputs to the amygdala promotes the formation of extinction memories [10]. We suggest the notion that enhanced transmission in this pathway may possibly reflect a core mechanism underlying angiotensin regulation of extinction learning. Angiotensin receptors are densely expressed in limbic and prefrontal regions critically engaged in extinction [16-18] and are considered to modulate learning-related neuroplasticity. LT is a selective competitive antagonist of the angiotensin II type 1 receptor but also increases availability of angiotensin II-converted angiotensin IV-an agonist at the AT4 receptor subtype. The AT4 system is thought to play a role in neuroplasticity and learning and memory [16, 17, 28, 51], a mechanism that is suggested to likely contribute to LT-induced extinction enhancement.

Consistent with previous animal models demonstrating the potential of LT to enhance extinction in rodents [19, 20], the present study successfully demonstrated the potential of a single, low-dose administration of LT to facilitate extinction learning in humans. In the context of recent findings suggesting a direct association between extinction-related vmPFC functioning and exposure therapy success [52], the current results indicate that LT represents a highly promising candidate to augment the efficacy of exposure-based interventions in therapeutic settings. On the neural level the effects of LT were mediated by circuits consistently involved in anxiety disorders with exaggerated threat reactivity and deficient extinction being associated with decreased vmPFC activation and dysfunction in the vmPFC-BLA circuit [6, 8, 11, 13]. Importantly, dysregulations in this circuitry normalize during the course of successful treatment [53] suggesting that they represent treatment-responsive elements rather than stable - markers and consequently promising targets for innovative therapeutic interventions. A previous human neuroimaging study reported that LT improves threat discrimination in high anxious individuals [18] and together with LT’s excellent safety record in clinical applications [22], the currently observed extinction enhancing potential and selective effects on vmPFC-BLA threat signaling, make this drug an attractive candidate for augmenting the effects of exposure therapy.

However, despite these initial promising results subsequent studies need to (1) examine effects of LT on subsequent extinction consolidation and recall in humans [as previously demonstrated in rodents 19], (2) determine the generalization of the effects to female subjects and, finally, (3) evaluate its potential to enhance exposure-based interventions in clinical trials.

Overall, the present results indicate an important regulatory role of the RA system in fear extinction learning in humans that are mediated by modulatory effects on vmPFC threat processing and its interaction with the amygdala. From a clinical perspective adjunct LT-treatment may represent an innovative strategy to enhance the efficacy of exposure-based interventions.

## Acknowledgements

We would like to thank Marianne Cumella Reddan for assistance with the threat expression analysis. This work was supported by grants from National Natural Science Foundation of China (NSFC) [91632117; 31530032]; Fundamental Research Funds for the Central Universities [ZYGX2015Z002]; Science, Innovation and Technology Department of the Sichuan Province [2018JY0001].

## Author contributions

F.Z. and B.B. designed the study, analyzed the data and wrote the manuscript. F.Z., Y.G., F.X., J.L., P.F., C.L, and W.Z. conducted the experiment. P.F., T.F., A.G., R.E. and K.K. revised the manuscript draft.

### Data and materials availability

Unthresholded group-level statistical maps are available on NeuroVault (https://neurovault.org/collections/4722/) and code that supports the findings of this study is available from the corresponding author upon reasonable request.

## Supplementary Methods

### Participants

We recruited 70 healthy male university students in the present study. Exclusion criteria included color blindness; systolic/diastolic blood pressure > 130/90 mmHg or < 90/60 mmHg; Body Mass Index (BMI) > 30 kg m^-2^ or < 18 kg m^-2^; current or regular substance or medication use; current or history of medical or psychiatric disorders; any endocrinological abnormalities or contraindications for LT administration and MRI. In line with previous studies targeting extinction processes [1] n = 8 participants (LT group, n = 3) who failed to acquire a conditioned threat response (average CS^+^ SCR < CS^-^ SCR during acquisition) were excluded. Three additional participants (LT group, n = 2) were excluded due to incomplete SCR data (technical issues). During acquisition of the primary neural outcome assessment (extinction) no subjects showed excessive head motion (> 3 mm translation or 3°rotation) leading to a final sample of n = 30 LT- and n = 29 PLC-treated subjects for the evaluation of the primary hypotheses of the present study. One participant (LT) was excluded from the fMRI analyses of the acquisition phase due to excessive head motion during acquisition. The sample size is in line with previous pharmaco-fMRI studies examining the extinction enhancing potential of pharmacological agents (n = 31 per group [2]) and previous studies examining the effects of LT on memory (n = 15 per group [3]) and fear-associated brain activity (n = 15 per group [4], see also sample size and power calculation in Reinecke, et al. [4]).

### Procedure

Losartan crosses the blood-brain barrier [5, 6], and following oral administration peak plasma levels are reached after 90 minutes [7, 8]. Whereas effects on cardiovascular indices only become apparent after 3 hours, effects at central receptors following intravenous administration have been observed after 30 minutes [6]. In line with the pharmacodynamic profile of LT [7, 9] and previous studies examining the cognitive enhancing properties of LT in healthy subjects [3] the experimental paradigm (extinction) started 90 minutes after administration. Although previous studies reported a lack of effects of single-dose LT administration on cardiovascular activity 3h after treatment [7], blood pressure and heart rate were assessed before drug administration, as well as before and after the extinction paradigm to further control for potential confounding effects of LT on cardiovascular activity (**Fig. 1**). Throughout the experiment affective state (Spielberger State–Trait Anxiety Inventory (STAI) and the Positive and Negative Affective Scale (PANAS) before drug administration and after the extinction paradigm [10, 11]) were acquired to control for unspecific emotional effects of LT (**Fig. 1**).

During acquisition participants underwent an adapted version of a validated Pavlovian discrimination threat conditioning procedure with partial reinforcement. Briefly, one colored square (CS^+^, 4s) coincided with a mild electric shock (US, 2ms) to the right wrist with 43% contingency, whereas the other differentially colored square (CS^-^, 4s) was never paired with the US. Acquisition included two runs and each run contained eight non-reinforced presentations of the CS^+^ and the CS^-^, intermixed with an additional six presentations of the CS^+^ paired with the shock (CS^+^U). In line with Schiller, et al. [12] first trial during acquisition was a reinforced one and we included one occurrence of consecutive reinforced CS^+^U trials. Stimuli were presented in a pseudorandom order with a 9-12s interstimulus interval (ISI) (fixation-cross) that served as low level baseline. Before the experiment participants were instructed to pay attention to the stimuli and find out the relationship between the stimuli and the shocks. In post-acquisition interviews, all of the subjects reported correct relationships.

The extinction procedure consisted 2 runs with each run encompassing 10 CS^+^ trials without the US and 10 trails of the CS^-^. The colors of stimuli, durations of trials and the range of ISI were identical to the acquisition phase. No trial-type was repeated more than two times in a row during either acquisition or extinction. The stimulus presentation and timing were optimized employing the make_random_timing.py script implemented in AFNI (Analysis of Functional NeuroImages, https://afni.nimh.nih.gov/) and were selected from 10,000 potential designs to maximize design efficiency. Two color sets were used and balanced across subjects to control for effects of the designated colors (color set A: red = CS^+^, blue = CS^-^; color set B: Green = CS^+^, pink = CS^-^).

### Unconditioned Stimulus and Physiologic Measurement

A mild electric shock (2ms duration) generated by a Biopac stimulator module STM100C and a STIMSOC adapter (Biopac Systems, Inc.) served as US. Shocks were delivered via two MRI-compatible electrolyte gel supported Ag/AgCl electrodes attached to the subject’s right wrist. Immediately before the start of the experiment, shock intensity levels were adjusted on an individual level by delivering gradually increasing shocks until the shock reached the level that participants reported as “highly uncomfortable, but not painful”. SCRs were assessed at a sampling rate of 1kHz from pre-gelled Ag/AgCl laminated carbon snap electrodes (EL508, Biopac Systems, Inc.) attached to the first and second fingers of the left hand between the first and second phalanges via Biopac Module EDA100C-MRI module attached to a MP150 (Biopac Systems Inc.).

### Skin Conductance Response Analysis

The level of a skin conductance response per se was defined as the maximum of the low-pass filtered (0.1Hz) conductance signal during a time window beginning 1 second and ending 5 seconds after stimulus onset minus baseline (the mean conductance in the 2s immediately before the onset of the stimulus). Responses below 0.02μS were encoded as zero [13-15]. To account for individual variability, the raw SCR scores were square root transformed and range corrected by dividing each response by the mean square root transformed US response during initial acquisition[15, 16]. Following previous procedures [13, 15, 16], we divided both CS^+^ and CS^-^ stimuli of each run into early (first half) and late (last half) phases and the mean SCR for CS^-^ stimulus was subtracted as baseline from CS^+^. The differential SCR response (CS^+^ - CS^-^) was regarded as the psychophysiological threat response to the conditioned fear stimulus. To initially test whether the threat responses were equally acquired to the conditioned threat between LT and PLC groups, we performed two-sample t-tests on the mean psychophysiological threat responses during acquisition. Note that we only included the non-reinforced trials, meaning that the CS^+^U trials followed by shocks were excluded from SCR analyses. To initially examine whether LT had unspecific effects on the SCR signal two-sample t-tests on the mean SCR for CS^-^ trials during extinction were conducted. To test whether LT had an effect on extinction learning, a 2 × 2 × 2 mixed-design analysis of variance (mixed ANOVA) model with the within-subject factors “phase” (early, late) and “run” (run1, run2), the between-subject variable “treatment” (LT, PLC) and the mean differential SCR during extinction as dependent variable was performed. All statistical analyses were performed in SPSS 22.0 (SPSS Inc) with Greenhouse-Geisser correction for non-sphericity if indicated. Significant main or interaction effects were further disentangled using appropriate post hoc t-tests focused on differences between the treatment groups (Bonferroni corrected).

### MRI Data Acquisition and Preprocessing

Functional MRI data was acquired using a T2*-weighted echo-planar imaging (EPI) pulse sequence (33 transverse slices, repetition time = 2s, echo time = 30ms, slice thickness = 3mm, gap = 0.6mm, field of view = 200 ×200mm, resolution = 64 ×64, flip angle = 90°, voxel size = 3.1 × 3.1 × 3.6mm). To improve spatial normalization and exclude participants with apparent brain pathologies a high-resolution T1-weighted image was acquired using a magnetization-prepared rapid gradient echo (MPRAGE) sequence (176 sagittal slices, repetition time = 1900ms, echo time = 2.52ms, slice thickness = 1mm, field of view = 256 × 256mm, acquisition matrix = 256 × 256, flip angle = 9°, voxel size = 1 × 1 × 1mm).

Functional MRI data was preprocessed using SPM12. The first five volumes of each run were discarded to allow MRI T1 equilibration. The remaining volumes were spatially realigned to the first volume and unwarped to correct for nonlinear distortions possibly related to head motion or magnetic field inhomogeneity, co-registered to the structural image, normalized to the Montreal Neurological Institute (MNI) space using a two-step procedure implementing segmentation of the T1-weighted image and application of the resulting deformation parameters to the functional images (interpolated to 3 × 3 × 3mm voxel size), and spatially smoothed using an 8–mm full-width at half maximum gaussian kernel. For the multivariate voxel pattern analysis unsmoothed data was used.

### Analysis of fMRI Data

A two-level random effects general linear model (GLM) analysis was conducted on the fMRI signal for statistical analyses. For the threat acquisition phase, the first-level model included three boxcar regressors, one for each stimulus type: reinforced CS^+^, non-reinforced CS^+^ and CS^-^. For the extinction phase, separate boxcar regressors for each stimulus type (CS^+^ and CS^-^) were defined at 4 stages: early run1 (first half of run1), late run1 (second half of run1), early run2 (first half of run2) and late run2 (second half of run2). Each regressor was convolved with a canonical hemodynamic response function with its time derivative to reduce unexplained noise and better fit for the model. The fixation cross epoch was used as an implicit baseline, and a high-pass filter of 128 seconds was applied to remove low frequency drifts. Other regressors of non-interest (nuisance variables) included head motion parameters (Friston 24-parameter model [17]), and indicator vectors for spikes in head movements identified based on frame-wise displacement (FD) > 0.5mm [18]. Single-subject contrast images were obtained and then modeled at the second-level random effects analysis. For the acquisition phase the following contrasts were examined: (1) overall non-reinforced CS^+^ versus CS^-^ condition, to examine the threat acquisition networks across all participants, and (2) group differences on the contrasts of non-reinforced CS^+^ versus CS^-^ condition to control for pre-treatment group differences during threat acquisition.

### Leave-one-subject-out (LOSO) Cross-validation (CV)

We employed a LOSO-CV procedure [19] to examine the robustness of the LT effect. Briefly, for every participant the same group-level analyses were performed excluding the participant’s data and the resultant clusters showing the significant interaction effect were used to obtain subject-specific masks, thus the remaining subjects served as an independent localizer for the subject left out. Next, individual mean beta values were extracted from corresponding contrasts within the subject-specific mask for each participant separately and appropriate t-tests were performed. This procedure ensured that the data used to define the subject-specific masks were independent from the data used for the group comparison.

### Construction of an Independent Anatomical-vmPFC Region of Interest (ROI)

The anatomical vmPFC was created by combining the subcallosal cortex, frontal medial cortex, anterior cingulate cortex and paracingulate gyrus from the Harvard-Oxford Atlas (thresholded at 25% probability) and only contained regions inferior to the genu of the corpus callosum and medial to x = 20 and x = −20 [20]. To increase regional-specificity a vmPFC region of interest (ROI) was constructed independent of the neural activity patterns in the present study by centering a 9mm-radius sphere around the centroid coordinates (MNI_xyz_ = [0, 34, −15]) of the anatomical-defined vmPFC mask. Note that we also thresholded the atlas at 0% probability and the resultant centroid coordinates were similar (MNI_xyz_ = [0, 34, −16]) and the corresponding findings remained the same.

### Exploratory Single Trial Analysis

Based on our hypothesis that LT would accelerate extinction leaning trial-wise vmPFC activations during CS^+^ presentation were explored. To this end a GLM design matrix with separate regressors for each trial was constructed with each aligned to either the onset of the CS^+^ or the CS^-^ stimuli. Considering that single trial estimates are highly sensitive to movement artifacts and noise that occur during one trial, we excluded trials with variance inflation factor > 4 or grand mean β estimate > 3 or < 3 standard deviation (SD) from the grand mean [21, 22]. One subject (PLC) was excluded from the single trial analysis given the high number of excluded trials (n = 14) with > 3 SD from the mean. Number of excluded trials for the remaining subjects was small and did not significantly differ between the treatment groups (LT, mean ± SD = 3.233 ± 1.942; PLC, mean ± SD = 2.536 ± 1.732, *P* = 0.155). To compute the mean beta values in the vmPFC ROI for each threat stimulus, the mean beta values in the same region across all of the CS^-^ trials were subtracted away from each CS^+^ trial as baseline for each run separately. Next the resultant beta values for each threat stimulus (across the two runs) were subjected to trial-specific two-sample t-tests. To control for the multiple comparisons, the *P* values were further false discovery rate (FDR) corrected. Due to limited reliability of the noisy single-trial SCRs (e.g., easily affected by stimulus unrelated movement and concurrent breathing) [23], we did not perform single trial SCR analyses.

### Multi-Voxel Pattern Analysis (MVPA)

Following Reddan, et al. [24], we used a linear support vector machine (C = 1) implemented in Canlab core tools (https://github.com/canlab/CanlabCore, based on the Spider toolbox) to develop a multivariate pattern classifier for threat (CS^+^) and non-threat (CS^-^). The pattern classifier was trained on subject-wise univariate non-reinforced CS^+^ > baseline and CS^-^ > baseline contrasts during threat acquisition. To avoid overfitting, the classification performance was evaluated by a LOSO-CV procedure. Next we used the pattern expression values, generated by taking the dot-product between the whole-brain unthresholded classifier weights with participant-specific brain activity maps, to evaluate effects of LT-treatment during the early extinction phase with a two-sample t-test. To avoid circularity, the pattern expression estimates for each individual were obtained from patterns trained on other participants’ data. In line with a previous study [24] the pattern expression was designed to differentiate CS^+^ versus CS^-^ and was thus considered as a measure of neural threat response.

### Mediation Analyses

Mediation analysis tests whether the observed relationship between an independent variable (X) and a dependent variable (Y) could be explained by a third variable (M). Significant mediation is obtained when inclusion of M in a path model of the effect of X on Y significantly alters the slope of the X–Y relationship. That is, the difference between total (path c) and direct (non-mediated, path c′) effects of X on Y (i.e., c - c′), which could be performed by testing the significance of the product of the path coefficients of path a × b, is statistically significant. We performed voxel-wise mediation analyses to determine whether brain activation mediates the relationship between treatment and the psychophysiological threat responses. To this end, treatment effects on fMRI activation (path a) and fMRI correlates of psychophysiological threat responses controlled for treatment effect (path b), and the a × b mediation effect were explored on the voxel-level. Significance estimates were calculated at each voxel for each path as well as mediation effect through bootstrapping (10,000 replacements). Based on our *a priori* regional hypothesis and the key role of the vmPFC in extinction [15, 25-27], bootstrapped two-tailed *P*-values for each vmPFC voxel were corrected for multiple testing (small volume correction, SVC) using false discovery rate (FDR) correction.

In parallel to the voxel-wise mediation analysis, we employed an independent vmPFC-focused single mediation analysis to specifically explore the contribution of the vmPFC (independently defined ROI) partial threat expression (M, CS^+^ > CS^-^) to the treatment (X) induced acceleration of early extinction learning on the psychophysiological threat response (Y).

### Psychophysiological Interactions Analysis

To examine the effects of LT on the functional interplay of regions involved in extinction, a generalized psycho-physiological interaction (gPPI) analysis was conducted. Similar to previous studies [see e.g. ref. 28], we used a group-constrained subject-specific approach to define individual volume of interest (VOI) seed regions (9mm-radius, sphere centered at local maximum CS^+^ > CS^-^ activity in the early extinction within the significant vmPFC cluster showing LT effects). Four PLC participants didn’t show greater activation for CS^+^ stimulus relative to CS^-^ stimulus in any voxel within the cluster and so the VOIs were centered at the group-level maximum.

A two-sample t-test was then conducted using FSL’s Randomise Tool to examine LT-effects on vmPFC functional connectivity for the CS^+^ versus CS^-^ during early extinction. Based on our *a priori* hypothesis the voxel-wise FDR correction was adapted to the structurally-defined bilateral amygdala (Harvard-Oxford Subcortical Structural Atlas, thresholded at 25% probability).

## Supplementary Results

### Acquisition Results

During the pre-treatment acquisition phase, both groups exhibited CS discrimination with enhanced SCR to the CS^+^ relative to the CS^-^ (**Fig. 1b**), confirming successful threat acquisition. On the neural level acquisition of threat was accompanied by stronger CS^+^ versus CS^-^ responses in the threat acquisition networks [see e.g. refs. 1, 29] (**supplementary Fig. 1**).

### Losartan Treatment Enhanced Threat-signal Specific vmPFC Activity

A more detailed examination of the treatment effects by means of extraction of the condition-specific neural signal from the independently (structurally) defined vmPFC ROI additionally revealed significant main effects of stimulus type (*F*_(1, 57)_ = 15.630, *P* < 0.001, partial η^2^ *=* 0.215, vmPFC activity was decreased to CS^+^ compared to CS^-^) and run (*F*_(1, 57)_ = 16.848, *P* < 0.001, partial η^2^ *=* 0.228 with increased vmPFC activity in run2 in relative to run1), as well as a significant treatment × stimulus type interaction effect (*F*_(1, 57)_ = 17.353, *P* < 0.001, partial η^2^ *=* 0.233) during early extinction. Post hoc comparisons between the treatment groups demonstrated that the interaction effect during early extinction was driven by a LT-induced selective increase of vmPFC responses to the CS^+^ (*t*_(57)_ = 2.777, *P* = 0.007, d = 0.723), in the absence of significant effects on the CS^-^ (*t*_(57)_ = −1.098, *P* = 0.277, d = −0.286). Post hoc comparisons between the stimulus types showed that in the PLC group the BOLD signal in response to the CS^+^ was decreased relative to the CS^-^ (*t*_(28)_ = −5.685, *P* < 0.001, d = −1.056). In contrast, LT-treated subjects exhibited a comparable vmPFC BOLD response to the CS^+^ compared to the CS^-^ (*t*_(29)_ = 0.152, *P* = 0.881, d = 0.028). In addition, in line with previous studies [30, 31] exploratory post-hoc paired sample t-tests showed conditioned threat responses with a decrease in BOLD signal for the CS^+^ relative to the CS^-^ in early extinction in PLC-treated subjects in both run1 (*t*_(28)_ = −3.917, *P* < 0.001, d = −0.727) and run2 (*t*_(28)_ = −4.829, *P* < 0.001, d = −0.897), reflecting persistence of the acquired conditioning memory. In contrast LT-treated subjects exhibited comparable responses to the CS^+^ and CS^-^ in both runs (*P*s > 0.6), suggesting that following LT-treatment differential vmPFC responsivity was already normalized during early extinction.

### MVPA Results

To test whether LT-treatment reduces neural threat response during early extinction, a whole-brain multivariate threat-predictive pattern was decoded [24]. Results showed a classification accuracy of 87.93% (*P* < 0.001) and sensitivity and specificity to threat were 0.862 (CI: 0.766-0.944) and 0.897 (CI: 0.811-0.967) respectively suggesting that the neural threat-predictive pattern could effectively distinguish neural threat versus non-threat processing (before treatment). When thresholded (bootstrapped 10,000 samples) and corrected for multiple comparisons (*q* < 0.05, FDR corrected), the neural threat-predictive pattern encompassed a distributed threat representation network including the vmPFC, insula, dorsal anterior cingulate cortex, thalamus and hippocampus resembling findings from mass-univariate analyses of threat acquisition in the present as well as previous studies [overview see ref. 29] (**supplementary Figs. 2a,b**).

Based on our *a priori* regional hypothesis and the key role of the vmPFC in the reduction of threat expression during extinction [15, 25-27] an additional analysis specifically focused on the effects of LT on the vmPFC partial threat expression. Briefly, the unthresholded neural threat-predictive pattern was masked by the anatomical vmPFC mask and next applied to early extinction activation maps with the higher partial pattern expression chosen as the threat stimulus (i.e., a forced-choice test). Results revealed that only the PLC group (accuracy = 89.66%, *P* < 0.001) but not the LT group (accuracy = 56.67%, *P* = 0.585) exhibited neural threat experience during early extinction.

### Mediation Results

A key question is how LT-treatment leads to accelerated extinction learning as reflected by reduced psychophysiological threat responses in early extinction. Based on previous studies indicating crucial contributions of the vmPFC to extinction and the facilitation of extinction [26, 27, 32-34] we hypothesized that treatment-induced effects on vmPFC activation would critically mediate accelerated extinction. We found that LT-treatment significantly contributed to both the reduced psychophysiological threat responses (path c; *t*_(57)_ = −2.179, *P* = 0.034) and enhanced vmPFC activation (path a; peak MNI_xyz_ = [6, 36, −21], Z = 4.616, *q* < 0.05, SVC-FDR corrected in pre-defined vmPFC mask, k = 544, **Figure 4A in yellow**). Furthermore, a significant negative correlation between vmPFC activity and the psychophysiological threat responses (path b; controlled for treatments, peak MNI_xyz_ = [-6, 48, −15], Z = −4.882, *q* < 0.05, SVC-FDR corrected, k = 221, **Figure 4A in green**) as well as a significant negative mediation effect in the vmPFC (peak MNI_xyz_ = [-3, 45, −18], Z = −3.795, *q* < 0.05, SVC-FDR corrected, k = 139, **Figure 4A in blue**) were observed. Importantly, a vmPFC cluster exhibited conjunction effects of both LT-treatment (path a) and fear expression (path b) as well as a mediation effect (peak MNI_xyz_ = [-3, 45, −15], Z = −3.692, *q* < 0.05, SVC-FDR corrected, k = 138), suggesting that activation in this region significantly mediated LT-induced attenuation of psychophysiological threat responses.

In parallel, an independent vmPFC-focused (ROI) mediation analysis demonstrated that LT-treatment significantly reduced the vmPFC partial threat expression (path a, b = −0.007, Z = −3.41, *P* < 0.001) and that lower vmPFC neural threat expression led to reduced psychophysiological threat expression with treatment as an adjustor (path b, b = 9.511, Z = 4.323, *P* < 0.001). More important, a significant mediation effect was found (a × b, b = −0.066, Z = −3.715, *P* < 0.001). In summary, these results suggested that LT enhanced early extinction learning through vmPFC processing.

### Visualization

Statistical maps were visualized using the Connectome Workbench (https://www.humanconnectome.org/software/connectome-workbench) and Mango (http://ric.uthscsa.edu/mango/). Behavioral data were plotted using Seaborn (https://seaborn.pydata.org/) and Dabest (https://github.com/ACCLAB/DABEST-python) [35].

---

**Supplemental Table 1.**
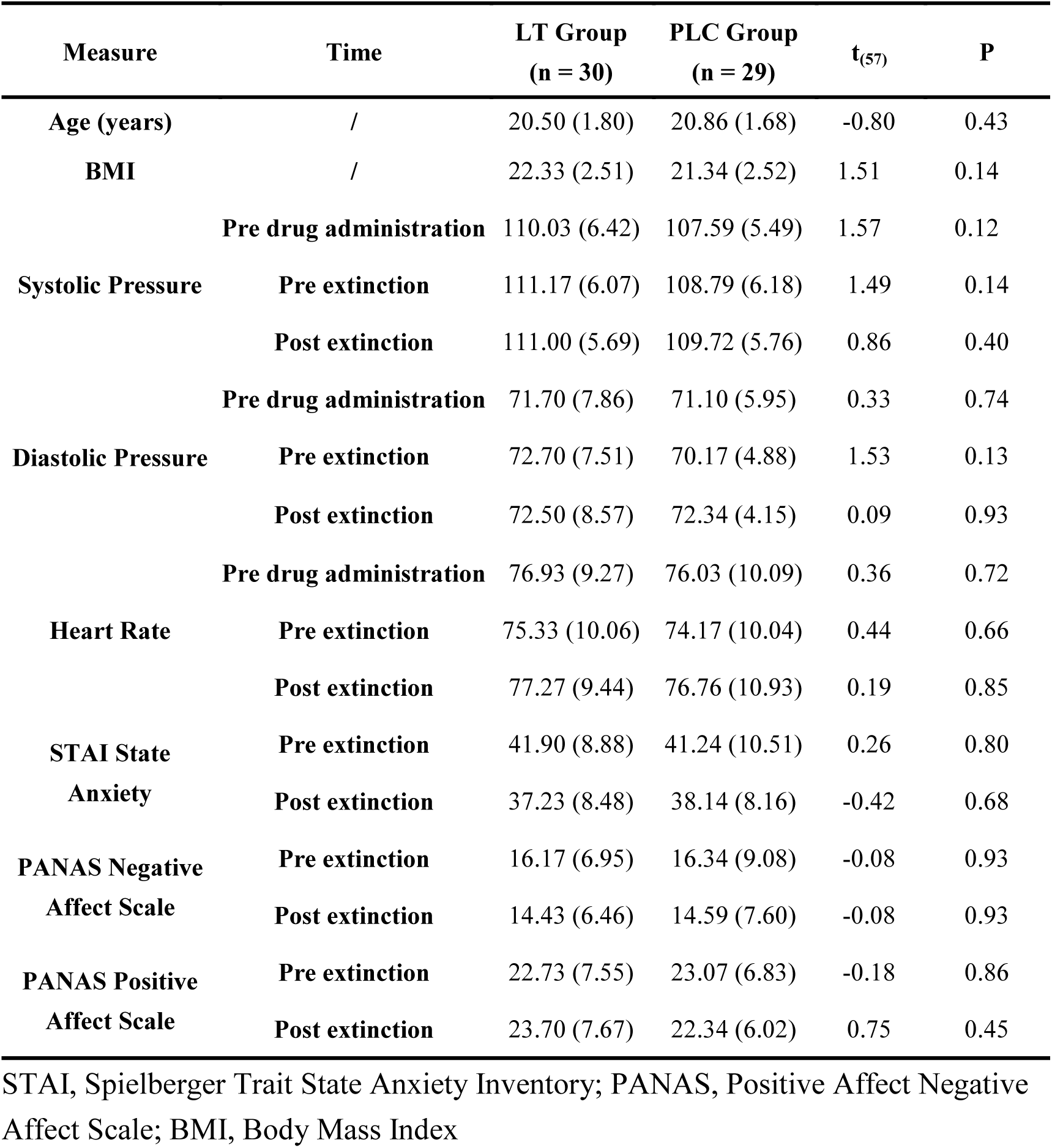
Participant demographics and neuropsychological performance STAI, Spielberger Trait State Anxiety Inventory; PANAS, Positive Affect Negative Affect Scale; BMI, Body Mass Index

## Supplemental Figures

**Supplemental Fig.1.**
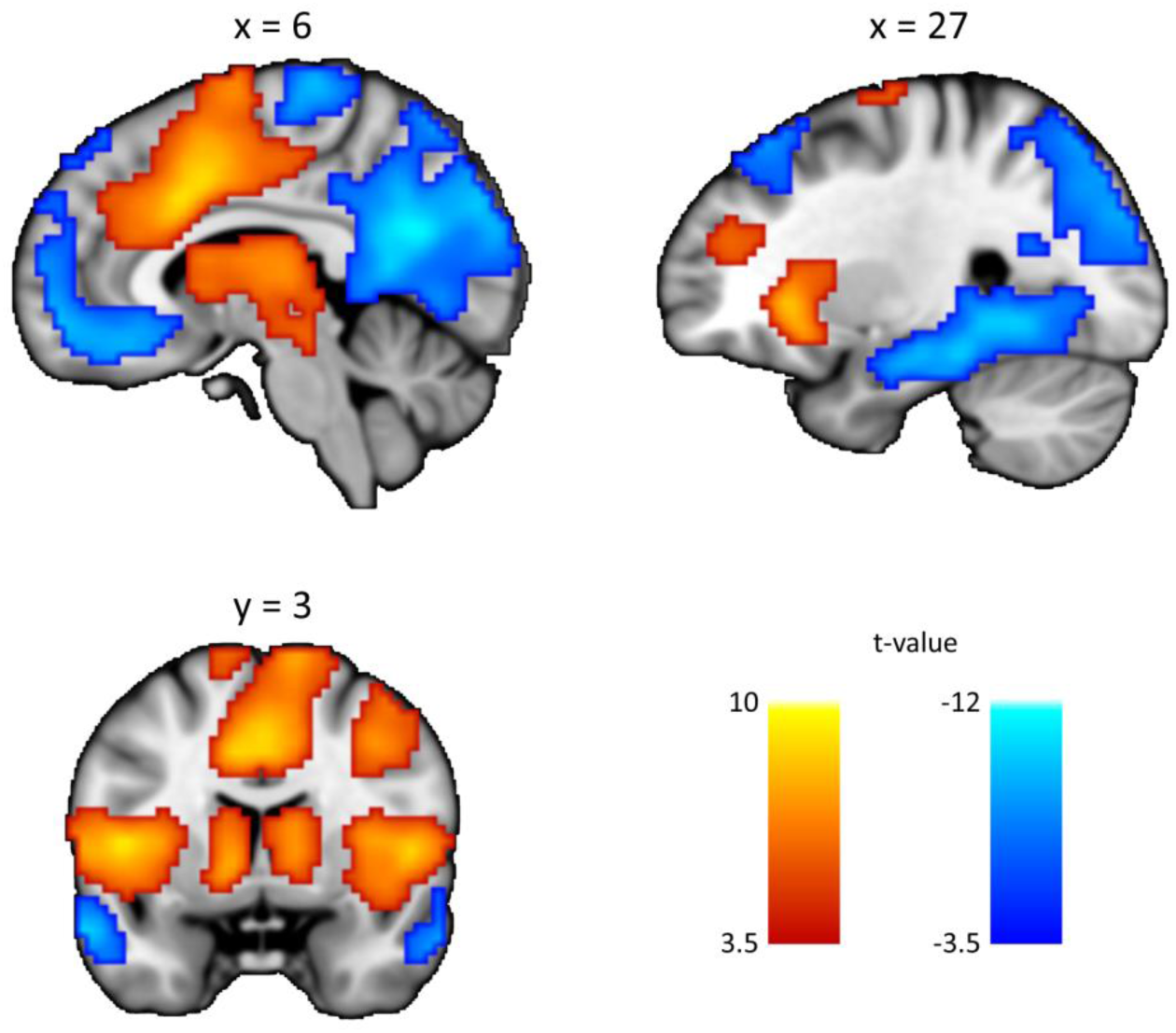
Non-reinforced CS^+^ versus CS^-^ BOLD responses during acquisition, across both experimental groups. Images displayed at *P* < 0.05, cluster-level family-wise error (FWE)-correction with a cluster-forming threshold of *P* < 0.001, two-tailed. Hot color indicates CS^+^ > CS^-^, whereas cold color indicates CS^+^ < CS^-^.

**Supplemental Fig.2.**
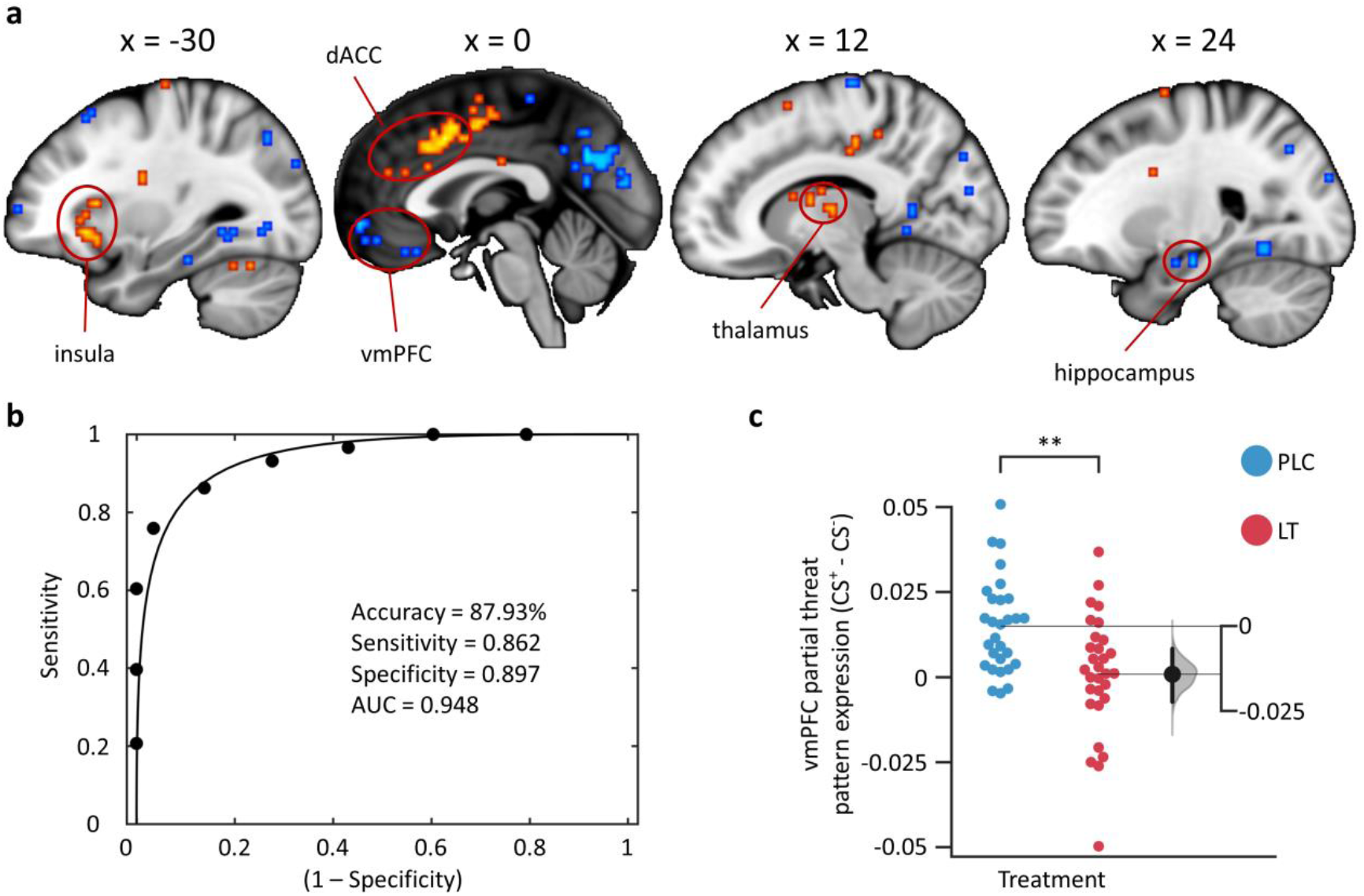
Multivariate neural threat-predictive pattern results. (a) Neural threat-predictive pattern, consisting of voxels in which activity reliably predicted threatening (unreinforced CS^+^) versus non-threatening (CS^-^) stimuli during threat acquisition. The map shows weights that exceed a threshold (*q* < 0.05, FDR corrected based on bootstrapped 10,000 samples) for display only. dACC, dorsal anterior cingulate cortex; vmPFC, ventromedial prefrontal cortex. Hot color indicates positive weights and cold color indicates negative weights. (b) ROC plot. The neural threat-predictive pattern yielded a classification accuracy of 87.93% in a leave-one-subject-out cross-validation (LOSO CV) procedure. (c) Losartan treatment reduced the partial threat pattern expression of the vmPFC. \*\**P* < 0.01. The filled curve indicates the null-hypothesis distribution of the difference of means (Δ) and the 95% confidence interval of Δ is illustrated by the black line.

